# Constancy and Change in the 5’UTR of Yellow Fever Virus

**DOI:** 10.1101/2020.07.17.207902

**Authors:** Stephen J. Seligman

## Abstract

The nucleotide composition of the 5’UTR of the yellow fever virus (YFV) has been reported to be the most constant sequence in the viral genome, but a comprehensive analysis of this constancy has not been presented. The current report is an analysis of the 5’UTRs from 48 sequences deposited in GenBank representing the seven described genotypes, five in Africa and two in the Americas. The YFV 5’UTRs consist of 118-120 nucleotides, 92% (110/119) of which are constant in all sequences. The constancy is impressive and suggests that many participate in significant viral functions. Remarkably, analysis of the non-constant nucleotides revealed that in some instances the non-constant nucleotide changes persisted in one or a restricted number of related genotypes and were from sequences isolated over a considerable span of years. This constant feature of non-constant nucleotides is consistent with the concept that the changes were in response to different environmental features such as changes in mosquito hosts or animal reservoirs, particularly as a consequence of spread of YFV from Africa to the New World. Constancy of 5’UTR in general may be helpful in distinction of viral species. Lastly, the presence of sequences of constant nucleotides greater than 19 nucleotides suggests regions of the 5’UTR that may be exploited for use as non-codon RNA as treatment and diagnostic agents in a variety of viral diseases.

**Importance:** The 5’UTR is arguably the most neglected portion of the viral genome. It is frequently incomplete in the sequences deposited as otherwise complete sequences in GenBank. The current report is an analysis of complete 5’UTR sequences selected from those deposited in GenBank and indicates that the 5’UTR is 92% conserved confirming that it is a highly conserved portion of the viral genome and suggesting that each conserved nucleotide may be functionally significant. Repeated occurrences of even non-constant nucleotides belong to a restricted number of genotypes raising the possibility that adaptation to new mosquito hosts and animal reservoirs such as those that accompanied spread of yellow fever virus from Africa to the Western Hemisphere are relevant. Knowledge of prolonged strings of invariable nucleotides in the 5’UTR has been used in designing a method for detecting YFV and may also be relevant for designing sequences for viral control of a variety of viruses.

## Introduction

YFV is the eponymous virus for the genus Flavivirus. Originally isolated in 1927 from a patient in what is now Ghana, the nucleotide sequence of the vaccine virus derived from YFV was reported in 1985 (1). YFV persists as the cause jungle yellow fever transmitted by various mosquito species (*Anopheles sp*. in Africa and by *Haemagogus* sp. and *Sabethes* sp. in the Americas). Although detailed observations have not been presented, non-human primates (NHPs) in Africa are thought to be resistant to fatal infection (2). In the Americas, some NHPs succumb to the virus (e.g. *Alouatta* sp. (howler monkeys) (3) and *Sapajus* sp (capuchins) (4), but others are usually resistant to lethal infection (e.g. *Callithrix* sp (marmosets) (5) and Leontopithecus sp (golden lion tamarins) (6)). In both Africa and the Americas, the feared complication of yellow fever is the development of urban yellow fever in which the virus is rapidly spread amongst humans by *Aedes aegypti* and is associated with a high human mortality rate.

YFV has a 5’UTR, a sequence (cds) encoding three structural and seven non-structural proteins, and a 3’UTR. The virus originated in Africa and spread to the Americas in the 17^th^ century in ships bearing slaves (7). Currently the virus persists in sub-Saharan Africa and in South America. Five genotypes are recognized in Africa and two in S. America (8). The genotypes are associated with different geographical regions. More recently sub-clade lineages of the S. American genotypes have become recognized (9). As with other viruses that lack a proof-reading mechanism in their RNA polymerases, mutations are common. Accordingly, considerable sequence diversity is anticipated.

The current report is an analysis of the YFV 5’UTR sequences deposited in GenBank. They were isolated from the seven genotypes (Table 1). Three aspects are evaluated: 1) Nucleotide constancy, 2) an analysis of constant features of non-constant nucleotides, and 3) suggestions are presented for further exploration. The results demonstrate that YFV 5’UTR consists of 118-120 highly conserved (92%) nucleotides and is much more conserved than nucleotides in the cds or the 3’UTR (10, 11), Seligman, S.J. unpublished).

**Table 1.**
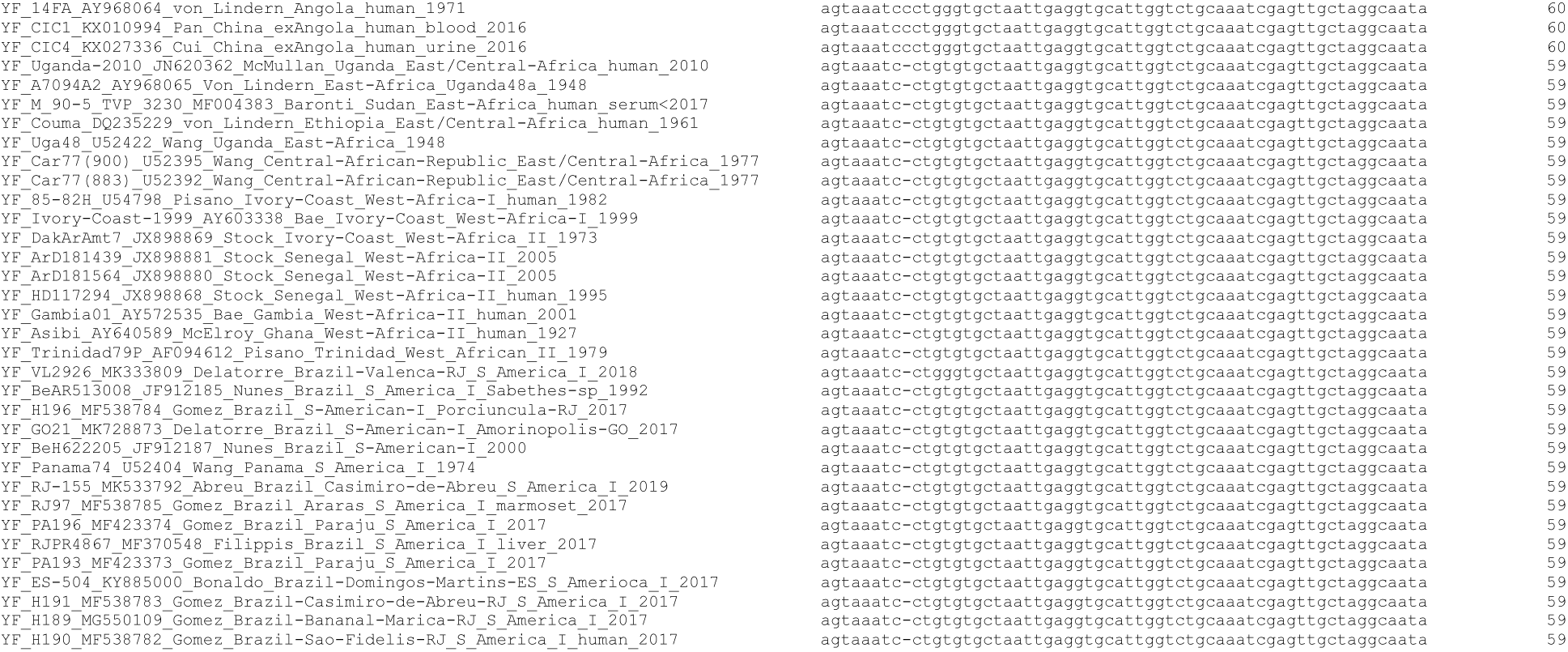
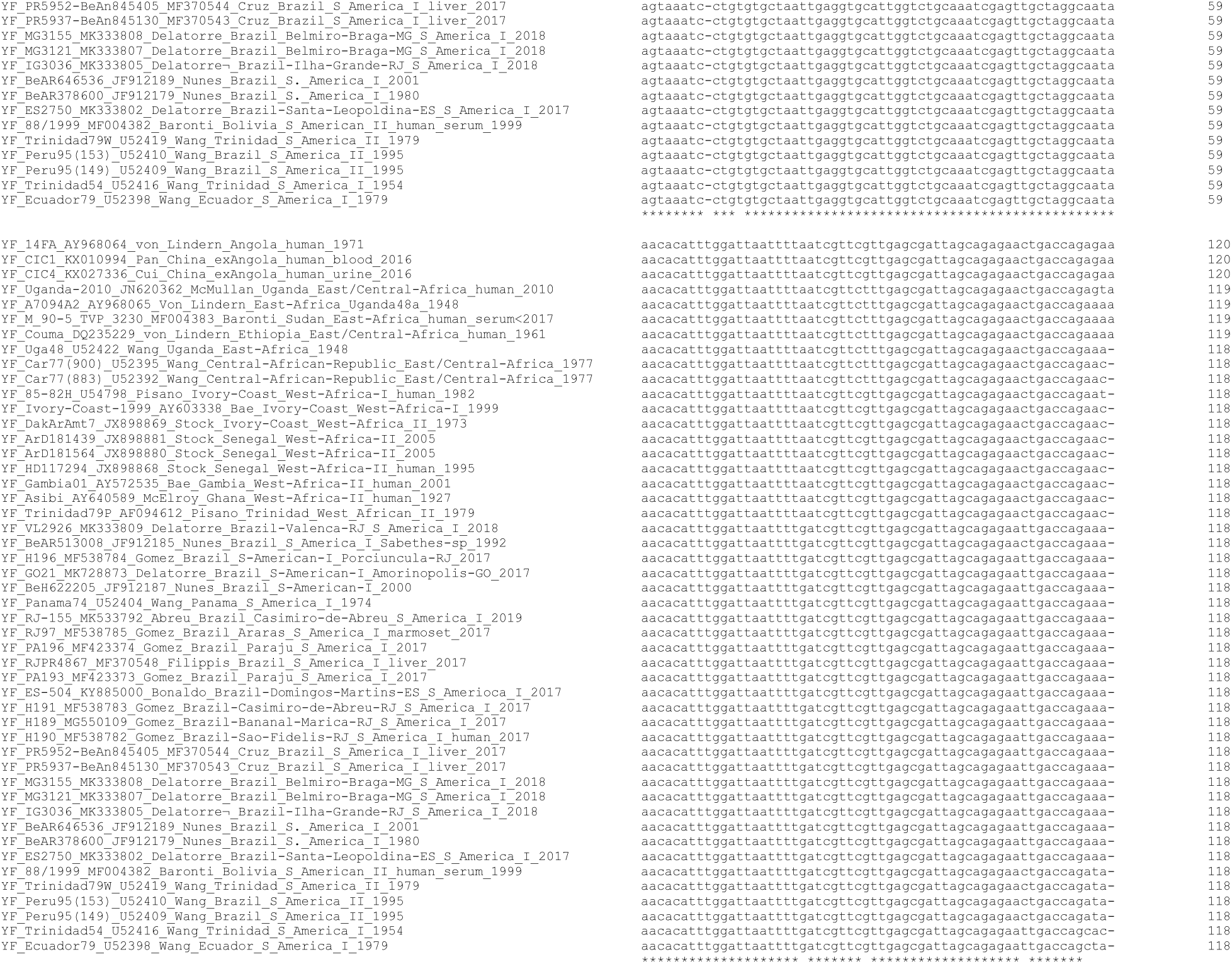
Alignment of yellow fever virus 5’UTR.

The relative constancy of YFV 5’UTR has been used in the development of a method to detect the YFV genome in point of care facilities and reference laboratories (11). Since YFV and other flaviviruses are noted for their nucleotide variability, the constant nucleotides in the 5’UTR suggest that each may have a raison d’etre. Analysis of some of the few non-constant YFV nucleotides reveals that they can persist for prolonged periods and be associated with a single genotype suggesting that there is a survival advantage for them as well to the mutation possibly related to different species of mosquito hosts and/or NHPs associated with particular geographic areas in which they propagate in South America (Table 2).

**Table 2.** Non-constant 5’UTR nucleotides in 48 yellow fever virus sequences.

| Nucleotide number | Nucleotide change | Genotypes with changed nucleotide |
| --- | --- | --- |
| 9 | a insertion | Angolan |
| 13 | g13t | Angolan |
| 81 | g81a | S. American I, Angolan |
| 89 | g89t | East Central Africa |
| 109 | c109t | S. American I |
| 117 | a117c | 2 sequences S. American I<br>From 1954 and 1971 |
| 118 | t118c | S. American II and 1 sequence from<br>S. American I |
| 119 | a119c | West Africa II, 2 sequences from<br>East/Central-Africa_1977, and 1 sequence from<br>S. America I 1954 |
| 119 | a119t | 1 sequence from East/Central-Africa_human_2010 and 1<br>sequence from West-Africa-I 1982 |
| 120 | a 120 deletion | An artifact of the alignment program. The deleted “a” is more reasonably considered to be the initial “a” in atg in the ensuing cds. |

To evaluate the extent to which constant yellow fever 5’UTR nucleotides remained constant in other flaviviruses, a similar analysis was done of 40 Zika virus 5’UTR sequences (data not shown). The Zika sequences contained 106-107 nucleotides, 90 of which were constant (84%).

In contrast with the non-constant 5’UTR YFV nucleotides, those of Zika virus were found in more recent isolates reflecting the explosive Zika epidemic. Whether the non-constant Zika nucleotides persist depends on future propagation of the virus.

The current analysis of constant and non-constant YFV 5’UTR nucleotides has the potential for yielding clues relating to the definition of virus species, the phenotypic implications of specific nucleotides, and the identification of stretches of conserved viral nucleotides helpful for the development of treatment non-coding RNAs.

## Methods

A search for suitable YFV sequences were made in GenBank, (https://www.ncbi.nlm.nih.gov/genbank/), an annotated collection of all publicly available DNA sequences. Search was made for sequences originating from different countries in Africa or Latin America in which yellow fever is or had been occurring. Sequences in which the 5’ end of the 5’UTR was incomplete were omitted. An alignment of 48 sequences was done. Aligned columns were evaluated in an effort to evaluate whether unique instances of non-constant nucleotides represented sequencing error or failure of these mutations to persist. Two such instances were found, a 1935 Brazilian isolate (U52389) and a Chinese isolate from a case acquired in Angola (KU921608) in 2016. In U52389 two different non-constant nucleotides and in KU921608 three occurred. A search using Blastn (https://blast.ncbi.nlm.nih.gov/) confirmed that the nucleotide changes in these two isolates were unique. The lack of repeat instances of these nucleotides in the relevant positions is consistent with either sequencing error or failure of the mutation to persist on subsequent viral replication thereby indicating that it is not an essential 5’UTR nucleotide, at least under the condition in which the virus was propagating.

## Results

Alignment of the 5’UTRs from 48 yellow fever strains including representatives from all of the seven recognized genotypes (8, 12, 13) indicates that their lengths varied from 118-120 nucleotides, 110 of which remained constant (92%) (Table 1). In contrast, 65% (6655/10230) of cds and 58% (266/459) of 3’UTR nucleotides were invariable (data not shown). In the case of cds variations, an increase in nucleotide constancy was noted with increasing size of the predicted viral protein (data not shown). The 3’UTR estimate is based on 22 isolates (14)). Sub-genomic stratification of repeated nucleotide sequence elements in the 3’UTR found in some YFV genotypes was not included in the calculations. Consequently more nuanced evaluations using sub-genomic sequences of the 3’UTR might increase the prevalence of constant nucleotides. The results do not necessarily establish all of the 19 or so consecutive conserved nucleotides necessary for viral growth inhibition by potential varieties of interfering RNA. The establishment of the combination of such nucleotides for the entire viral genome is beyond the scope of this manuscript.

Analysis of the non-constant nucleotides revealed two distinct patterns (Tables 1 and 2). In some instances (nucleotides 81, 89, 118, and 119), the non-constant nucleotides persisted for many decades in a single genotype suggesting a survival advantage of the relevant YFV in a particular geographic region that contained a specific mosquito species and/or non-human primate host. Instances of the occurrence of a given nucleotide change or changes in one or two sequences could be the result sequencing error. In any event failure of the nucleotide change to persist suggests minimal effect of their ability to confer a significant advantage to viral propagation.

## Conclusions

Analysis of constant and non-constant nucleotides in the 5’UTR of yellow fever virus that their persistence over the course of decades suggests that at least some nucleotides play a significant role in viral defense. Restraints in nucleotide changes in this region are likely to be of considerable interest. What mechanisms might be involved? In recent years considerable attention has been placed investigating the role of non-coding RNA in viral infections. Nucleotide sequences longer than 200 are usually considered as long non-coding RNA and have been found to be important both in host cell response and virus defense mechanisms (15, 16). In addition small non-coding RNA’s (RNAi) have a variety of formulations including both single-stranded RNA (micro (mi)RNA) (17) and double-stranded RNA (siRNA).Their actions may be promulgated either in *cis* or in *trans* (sfRNA) (18). Piwi-interacting (pi)RNA are found only in the testes and are involved in suppressing transposons (19).

Which, if any of these non-codon RNAs, are present in the 5’UTR is conjectural. Although 5’UTR nucleotides in alphaviruses have been implicated in host defense (20), the composition of RNAs in the 5’UTR that might be involved needs to be investigated systematically. Modern editing techniques changing selected 5’UTR nucleotides should be explored to investigate phenotypic changes in viral properties. Of particular interest is the possibility that nucleotide sequences from the 5’UTR are crucial in enabling YFV to propagate in primates and/or mosquitoes. Although no experiments have thus far reported systematically investigating the constancy of nucleotides in the 5’UTR, the effectiveness of the varieties of small interfering RNA in the control of virus infection is a field of active investigation (21-25). Should suitable conditions be found in which 5’UTR nucleotide sequences are determined to be inhibitory to YFV, parenteral delivery methods of the relevant inhibitory nucleotides would need to be developed to achieve a therapeutic effect. Most intriguing is the possibility that these concepts may also be applicable to other viruses such as coronaviruses, an idea of special significance in the control of rapidly emerging pandemics.

## Supplemental Material

None

## Acknowledgements

This work was initially stimulated by discussions with Charlie Rice

